# Enterovirus RNase L inhibiting RNAs are highly conserved with limited phylogenetic distribution

**DOI:** 10.64898/2026.06.29.735259

**Authors:** Samantha I. Zangari, Madeline E. Sherlock, Jeffrey S. Kieft

**Author notes:** To whom correspondence should be addressed: Jeffrey S. Kieft, New York Structural Biology Center 89 Convent Street, New York, NY, 10027.

## Abstract

RNA molecules form specific 3D structures that facilitate a variety of functions through interactions with other macromolecules. Many RNA viral genomes maintain these structures to interact with and evade host immunity machinery. One such element, the competitive inhibitor RNA (ciRNA), discovered in the protein coding region of the poliovirus serotype 1 (PV1) genome, inhibits a host antiviral protein, ribonuclease L (RNase L). Although some functionally essential structural motifs of the PV1 ciRNA have been studied, the extent of its evolutionary conservation and other structural requirements remained unexplored. Here we combined bioinformatic and biochemical techniques to further define the requirements of a functional ciRNA and assess its phylogenetic distribution. We systematically mutated ciRNA structural features, verifying that ciRNA inhibitory activity requires a conserved loop E motif and a long-range base-pairing interaction, but its peripheral stems are dispensable and in fact a circularly permuted version maintains function. A structure-based homology search identified potential ciRNAs across the *Picornaviridae* family, but only a subset of those tested were functional—all are in *Enterovirus coxsackiepol*. When structural features needed for function were transposed from PV1 ciRNA to an RNA unable to inhibit RNase L, the chimeric RNAs did not gain wild-type function, and chemical probing data revealed that these nonfunctional RNAs are unable to form the correct secondary structure. Overall, the dual constraints of encoding a protein and forming a specific functional structure appear to not only limit the sequence diversity, but also the phylogenetic distribution, of ciRNAs.

## Introduction

RNA is unique among biological macromolecules, serving as the template for protein production and performing many additional biological functions facilitated through its ability to form specific, unique, and complex 3D structures (Vicens and Kieft 2022). This characteristic of RNA is leveraged extensively by RNA viruses, which exploit the molecule’s multifunctionality to simultaneously confer important function to the genomic RNA while avoiding expansion of its complexity and size. Specifically, the same RNA molecule that encodes the viral proteins can contain elements that direct, regulate, or control important processes during viral infection such as translation, replication, or the production of new virions, among other functions (Bermudez et al. 2024). In addition, structured viral RNA elements often interact with cellular components to alter behavior in ways that promote infection such as disrupting host anti-viral responses (Steitz et al. 2011; Nagy and Pogany 2012; Jaafar and Kieft 2019; Filomatori et al. 2026).

Here, we focus an RNA element identified in the poliovirus serotype 1 (PV1) genome that possesses the surprising ability to inhibit the interferon-regulated endoribonuclease, RNase L (Han et al. 2007). RNase L is an important mammalian immune-response nuclease that plays an essential role in antiviral defense. Briefly, double-stranded RNAs produced during viral infection are recognized by oligoadenylate synthases (OASes), which catalyze the production of 2′-5′ linked oligoadenylates (2-5A) from ATP (Li et al. 2016). 2-5A binding to RNase L promotes dimerization and activates the enzyme (Dong et al. 1994; Carroll et al. 1997), which then preferentially cleaves both viral and cellular RNA substrates at UN/N dinucleotides, and at other sequences with lower efficiency (Floyd-Smith et al. 1981). Activated RNase L cleaves viral and host RNA to prevent translation of functional viral proteins and assembly of functional ribosomes. The resultant cleaved RNAs activate other antiviral responses, such as triggering apoptosis and interferon signaling to protect the host from infection (Baglioni et al. 1978; Chakrabarti et al. 2011). Structures of human and pig homodimeric RNase L reveal that one molecule of 2-5A contacts the N-terminal ankyrin-repeat of each monomer of RNase L, bridging the individual monomers (Han et al. 2014; Huang et al. 2014). The RNase L dimer is also stabilized by ATP or ADP, which binds to the pseudokinase domain and encourages important contacts in the center of the protein (Gusho et al. 2020). Lastly, structural studies of a catalytically inactive RNase L dimer bound to RNA substrate reveals an asymmetric RNA binding site within the RNase domains (Han et al. 2014).

The RNA element within the PV1 genome that inhibits RNase L activity spans a 225 nucleotide-long stretch of the single-stranded, positive-sense poliovirus genome, in the region that encodes the virus’s 3C protease (Han et al. 2007; Townsend et al. 2008a). The initial observation that led to the discovery of this element was that the PV1 genome is resistant to RNase L degradation, despite being rich in UN/N dinucleotides (Han et al. 2007). These studies revealed a specific structured RNA element in the genome that can inhibit RNase L outside the context of the PV genome. Specifically, in a minimal biochemical assay with activated RNase L, the 225 nt-long RNA was sufficient to inhibit the enzyme’s ability to cleave an ideal substrate. Subsequent kinetic analyses revealed that the RNase L-inhibiting RNA acted as a competitive inhibitor, directly competing with substrates for the RNase active site (Townsend et al. 2008a). This led to this RNA element being named the PV competitive inhibitor RNA (ciRNA; note that more recently this acronym has been used to refer to circular intronic RNA, an unrelated class of RNA). By interacting with and inhibiting activated RNase L, the PV1 ciRNA protects the viral genome, and potentially other RNAs, from degradation by the enzyme. Intriguingly, this inhibition is not permanent: RNase L activity is restored late in infection as virus assembly nears completion. This reactivation appears to be essential for the virus to spread, suggesting PV may ultimately co-opt the very pathway it initially suppresses to trigger cytopathic effects that facilitate release of assembled progeny (Han et al. 2007).

Although the active PV ciRNA sequence spans 225 nucleotides in the genome, the essential elements are discontinuous; that is, ~84 nucleotides can be removed from the middle of the sequence without losing the inhibitory function (nt 5823-5907) (Fig. 1A). The important regions form a secondary structure that includes multiple base-paired helices (P0-P3) connected by two three-way junctions (J1, J2) (Fig. 1A). P2 and P4 are capped by apical loops that are complementary to one another, forming base pairs in a tertiary “kissing loop” interaction that is important for full PV ciRNA function (Han et al. 2007; Townsend et al. 2008a; Townsend et al. 2008b; Keel et al. 2012a). In addition, one of the helical elements (P1) contains a loop E motif that is essential for function. Loop E motifs are a common type of RNA internal loop, often mediating RNA-RNA and RNA-protein interactions (Chastain and Tinoco 1991; Wimberly et al. 1993; Correll et al. 1997; Hampel and Burke 2001). The PV1 ciRNA structure appears to be conformationally dynamic, and although models for the overall folded architecture of this RNA have been proposed (Keel et al. 2012a), it is unclear whether the critical loop E and kissing loop motifs interact with one another, interact with RNase L directly, are required for proper tertiary RNA structure formation, or contribute to inhibition through a different mechanism.

**Figure 1.**
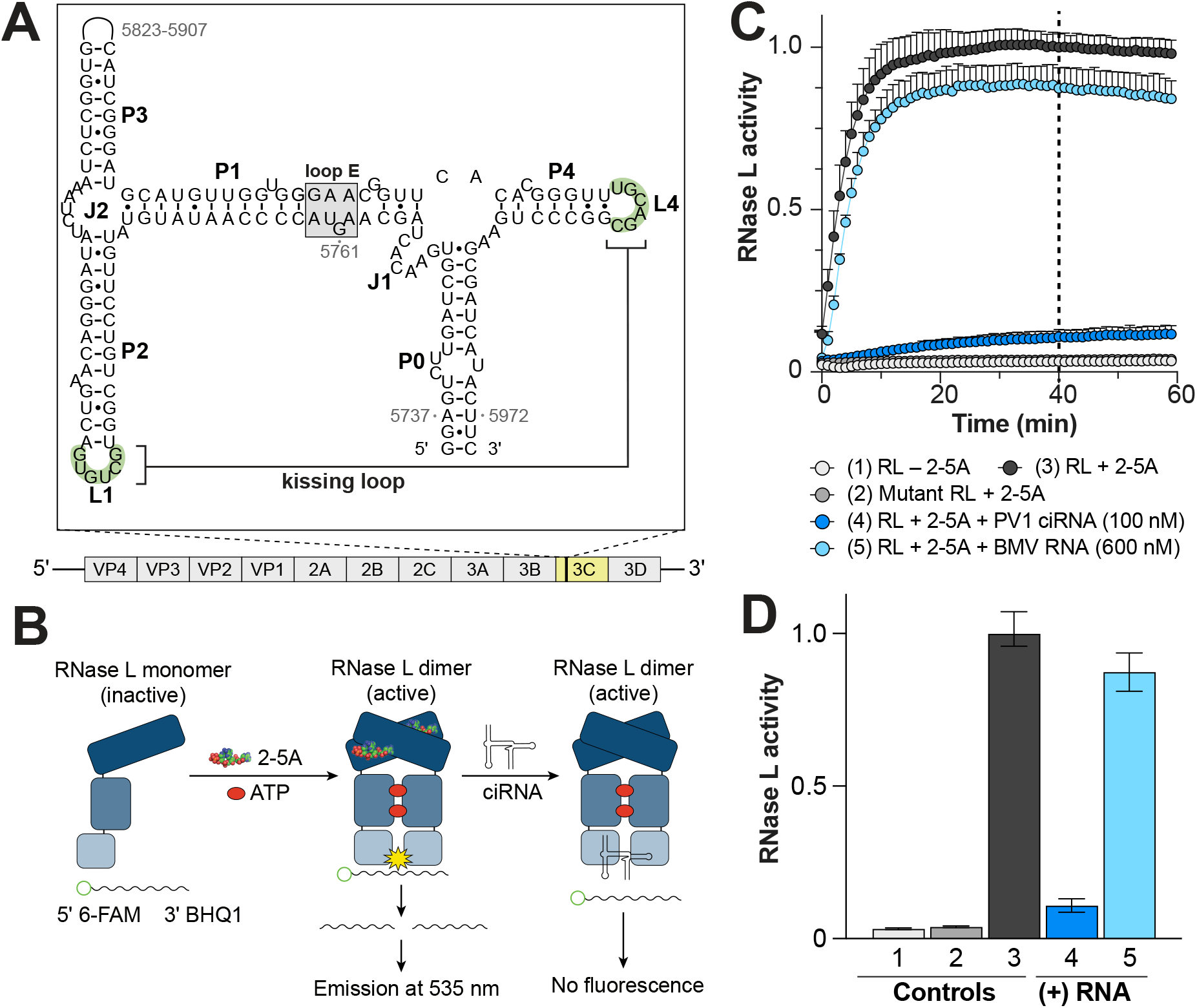
The PV1 ciRNA inhibits RNase L. **(A)** Secondary structure of PV1 ciRNA. The ciRNA is located in the 3C protease coding region of the PV1 genome. Paired elements (P), loops (L), and junctions (J) are labeled near each element. The long range base-pairing nucleotides of L1 and L4 (green) and loop E motif (gray) are highlighted. **(B)** Schematic of RNase L inhibition assay with various RNAs added. RNase L is activated with 2-5A and ATP and mixed with a fluorescently-tagged RNA probe and 0-600 nM ciRNA. Fluorescence is measured with and without a ciRNA to determine RNase L inhibiton. **(C)** Tiem course of the *in-vitro* assay to measure RNase L activity and inhibition of RNase L by PV1 ciRNA. Reactions containing RNase L (+/-2-5A) and probe RNA are measured at Ex/Em: 485/535 every minute over 1 hour at 25°C in triplicate (n=3). RNase L activity is normalized activated RNase L (+2-5A) control. **(D)** RNase L activity assay values at 40 min (n=3).

A ciRNA was first identified in PV1 and has been predicted to exist in few closely related coxsackieviruses (CVs) based on sequence conservation (Han et al. 2007), the full phylogenetic distribution remained unknown. Since the ciRNA’s requirements to serve as a template for protein synthesis constrains evolution of the inhibitory structure (and *vice versa*), it was plausible that other enteroviruses have a ciRNA within the 3C protease coding region. However, whether ciRNAs are broadly conserved across enteroviruses was unknown, as well as to what degree sequence variation occurs and is tolerated. To address this, we used in vitro reconstituted inhibition assays with the PV1 ciRNA to more fully determine functionally essential motifs, then used a secondary structure homology-based bioinformatic search to find other putative examples of ciRNAs. We tested the ability of these putative ciRNAs to inhibit RNase L and used chemical probing to determine the secondary structures of both inhibitory and non-inhibitory examples. Our results reveal strict primary, secondary, and tertiary structure requirements for ciRNA function within a defined RNA structure. The strict sequence and structural constraints on ciRNAs are mirrored by a limited phylogenetic distribution, with implications for RNA virus evolution and potentially therapeutic design.

## Results and Discussion

### Exploring the functionally essential components of a PV ciRNA

We used an established in vitro cleavage assay to measure the PV1 ciRNA’s ability to inhibit RNase L (Thakur et al. 2005; Han et al. 2007; Townsend et al. 2008a). In this assay, recombinantly expressed and purified human RNase L is activated with 2-5A and ATP, which then cleaves a single-stranded UA-rich probe RNA. This probe has a 6-FAM fluorophore and a black hole quencher (BHQ1) on its 5′ and 3′ ends, respectively. When this probe is cleaved, the BHQ1 no longer quenches the 6-FAM, resulting in emission at 535 nm (Fig. 1B). Emission values (RFUs) plotted as a function of time reveal the progression of target RNA cleavage by RNase L (Fig. 1C). Without ciRNA, RNase L cleaved most of the probe within the first 20-30 minutes. RFU values slowly decrease towards the end of the experiment, likely due to fluorophore bleaching (Fig. 1C). When PV1 ciRNA was added to the reaction at a concentration of 100 nM, the probe was cleaved at a significantly slower rate (Supplementary Fig. S1). This inhibition was specific to the ciRNA, as an unrelated but structured tRNA-like structure RNA from brome mosaic virus (BMV), of similar size to the PV1 ciRNA, did not inhibit RNAse L even at high concentrations of 600 nM (Fig. 1C&D, Supplementary Fig. S1). These results confirmed previously published data for measuring relative ciRNA inhibition of RNase L. For visual simplicity and to allow easy comparison between different ciRNA mutants, hereafter we use the fluorescence value at the 40-minute time point to report relative inhibition (Fig. 1D).

We used this assay to test the functional importance of several PV1 ciRNA secondary structure elements. First, we explored the length requirement of the P0 stem that forms by base pairing between the 5′ and 3′ ends, shortening it to 7 bp (d1) and 4 bp (d2) (Fig. 2A, Supplemental Fig. S2). Both mutant RNAs maintained their ability to inhibit RNase L similarly to wild type (Fig. 2B). Next, although it was previously shown that nucleotides 5823-5907 of the PV1 genome are not required for ciRNA function (Townsend et al. 2008a), the necessity of the P3 stem, which connects this intervening region to the ciRNA through junction J2, was not fully investigated. We therefore made two mutants, one with the P3 stem trimmed to 4 bp (d3) and the other with P3 completely removed, converting the J2 three-way junction to an internal loop (d4, Fig. 2A). Both d3 and d4 effectively inhibited RNase L (Fig. 2B), highlighting that P3 is not needed and thus a three-way junction formed between P1, P2, and P3 is not a necessary feature of a functional PV1 ciRNA.

**Figure 2.**
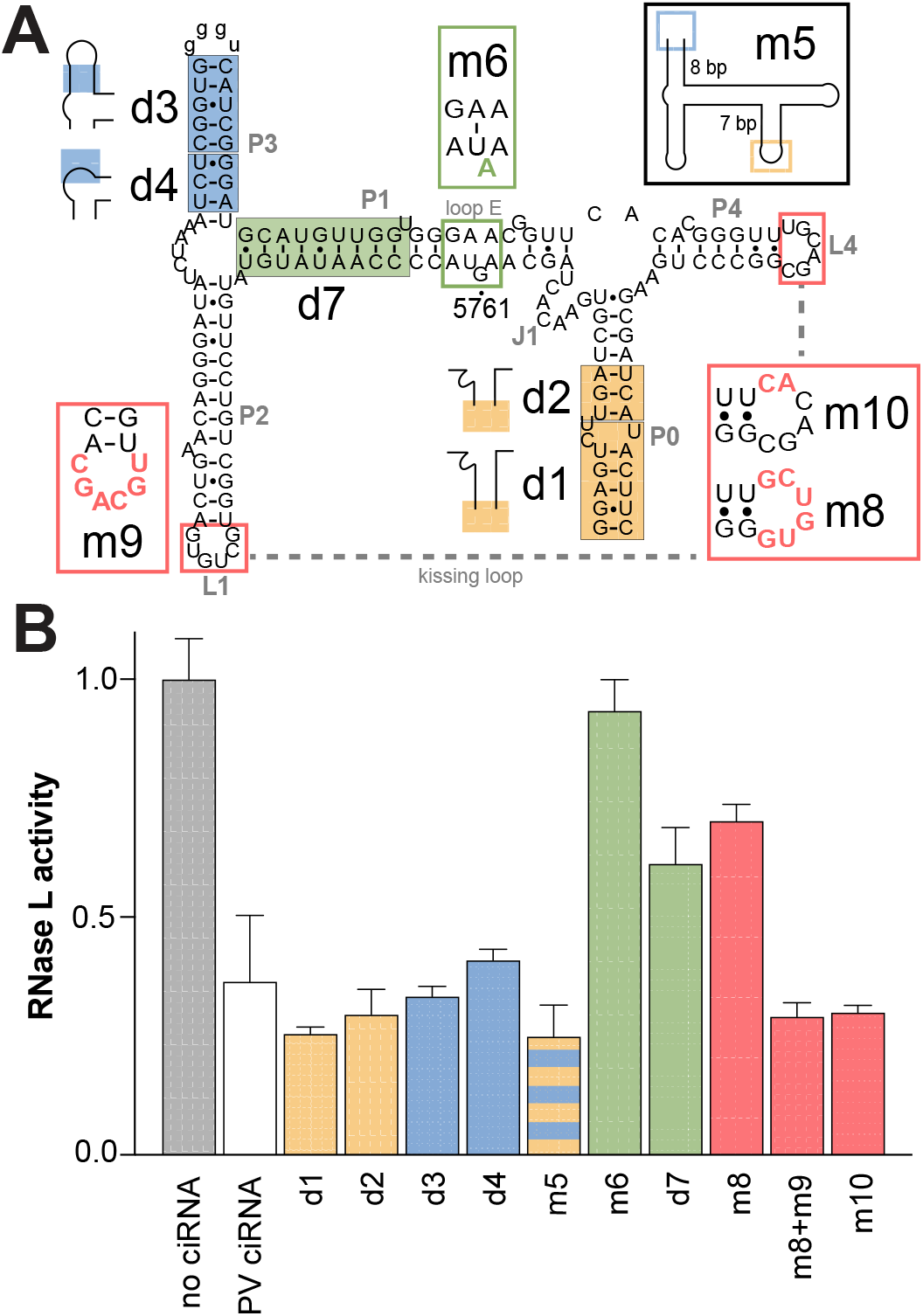
Mutational analysis of PV1 ciRNA. **(A)** Mutations (m, outlined) and deletions (d, shaded) plotted on ciRNA secondary structure; mutations in P0 (yellow), P1 (green), P3 (blue), and L1 and L4 (salmon). Full secondary structures of all mutants are contained in Supplementary Figure S2. **(B)** Results of RNase L inhibition assay for all mutants of panel A. Reactions containing ciRNA have 100 nM RNA. Replicates (n=3) plotted at 40 min.

Having shown that the PV1 ciRNA function does not depend on the sequence and length of the P0 and P3 stems, we hypothesized that a circularly permuted ciRNA would be functional. This ciRNA would have new 5′ and 3′ ends with the naturally occurring ends connected, resulting in an isomer capable of forming the same secondary structure as the original RNA (Pan and Uhlenbeck 1993). We generated and tested an RNA with new 5′ and 3′ ends in the P3 stem, and the original ends in P0 connected by a GNRA tetraloop (m5, Fig. 2A, Supplemental Fig. S2). This circularly permuted ciRNA inhibited RNase L as well as wild-type, consistent with the model of the ciRNA as a discrete functional unit. Its ability to inhibit the RNase depends on a specific secondary structure and presumably key internal tertiary interactions but is independent of surrounding RNA or primary sequence context.

We then explored the effects of mutations within the ciRNA designed to alter important tertiary motifs or long-range tertiary interactions. First, we altered the loop E motif with mutation G5671A (m6). Consistent with previous reports, alteration of the loop E motif by just a point mutation ablated ciRNA’s ability to inhibit RNase L (Townsend et al. 2008b; Keel et al. 2012a). Likewise, consistent with previous studies, targeting the L1-L4 kissing loop interaction by mutations that eliminate potential base pairing between them resulted in a substantial loss of function (m8, Fig. 2B). When both L1 and L4 were mutated simultaneously to restore potential base pairing (i.e., L1 and L4 sequences swapped, m8+m9), ciRNA function was fully restored. Finally, as the kissing loop interaction may contain two G-U pairs, we mutated two nucleotides in L4 to convert these to two G-C pairs, potentially increasing the stability of the long-range interaction. This mutation showed wild-type inhibitory function, but not increased function (m10, Fig. 2B). Overall, L1-L4 base pairing is less essential to ciRNA function as is the loop E motif but is still important for maximal ciRNA inhibition.

As the loop E motif and L1-L4 pairing are both important for full activity, we tested if their relative context within the overall ciRNA architecture is important. To do this, we shortened the P1 stem by 9 bps, or approximately one helical turn (d7; this had a the shorter P3 stem). Shortening of P1 reduced the ability of the ciRNA to inhibit RNase L, although it retained more inhibitory function than the mutant that directly disrupts the loop E motif (m6). While it is possible that the P1 primary sequence is important, a more likely explanation is that shortening P1 alters the orientation and spacing between the L1 and L4 loops, preventing their base pairing and thus negatively impacting ciRNA function. Consistent with this, shortening P1 in d7 and disruption to the kissing loops interaction (m8) resulted in a qualitatively similar effect on ciRNA function. Overall, these results show that the loop E and potential for L1-L4 base pairing are necessary but not sufficient for full function, and their context within a specific architecture is important

### Homology search to identify and catalog new putative ciRNAs

We used our enhanced understanding of PV1’s ciRNA to find and classify different putative examples of this element using a secondary structure-based homology search. We used the secondary structure of the 225-nt ciRNA sequences from poliovirus serotypes PV1 (NC_002058.3), PV2 (M12197.1), and PV3 (K01392.1) to create an initial seed alignment (Supplementary Fig. S3). These sequences were aligned by primary sequence and predicted secondary structure, ignoring putative tertiary interactions or long-range base-pairing. We used this alignment with the Infernal program to search the sequence database of all fully sequenced genomes within the *Picornaviridae* family, to which PV1 belongs (Nawrocki and Eddy 2013). This search yielded 317 candidate ciRNAs from 12 different viral genera, with over 98% of results belonging to the subfamily, *Ensavirinae* (Zell et al. 2021).

Since Infernal uses both secondary structure and primary sequence to search for other conserved structures, the resulting RNAs from this search are constrained by the protein-coding requirement of the ciRNA element. The sequences identified by this search were found within the annotated predicted 3C protease coding region of picornavirus genomes. As expected, the sequence diversity between candidate RNAs is primarily localized to wobble-position residues, and the highest primary sequence conservation is localized to regions encoding the catalytic residues of the 3C protease (Supplementary Alignment File). Interestingly, these catalytic residues are encoded within the region connected to the ciRNA by P3 and dispensable for inhibition function. Outside of the catalytic center, the 3C protease sequence is somewhat more variable among enteroviruses (Laitinen et al. 2016), and this likely allows for some variation to arise and for the evolution of a functional RNA structure to perform a noncoding function.

The primary sequences of the RNAs found in this search are insufficient to determine their ability to inhibit RNase L. Therefore, we chose several RNAs from this alignment to test for the ability to inhibit RNase L in the in vitro assay. Based on visual examination, several RNAs were chosen that contain a loop E motif and the potential to form a kissing loop interaction, others were chosen based on factors like primary sequence similarity to PV ciRNA and E-value, a metric for statistical significance of each identified sequence compared to the PV input sequences (a lower E-value is more significant) (Supplementary Fig. S4B). Not surprisingly, there is a correlation between percent sequence identity to poliovirus and a more significant E-value, again due to the constraints imposed by the protein sequence (Supplemental Fig. S4A). Additionally, the candidates chosen for validation mostly belong to *Enterovirus coxsackiepol* species but with other candidates from other enteroviruses. The AnaV-A1 RNA was chosen as an outlier because it does not belong to the same genus as the rest of the RNAs and it does not infect a mammalian host, unlike the other viruses.

The selected RNAs were in vitro transcribed and purified, then tested for function using the in vitro inhibition assay described previously. Interestingly, only RNAs from viruses belonging to *Enterovirus coxsackiepol* were able to inhibit RNase L, specifically CVA11, CVA13, CVA20, CVA21, CVA24, and EV-C99 (Fig. 3A). Sequences from CVA19, CVA22, and EV-C104, which are all members of the same viral species, did not show the ability to inhibit RNase L. Of the RNAs tested, only those with E-values below 1E-60 contain a functional ciRNA, as determined by our in vitro assay (Fig. 3B). Strikingly, the ability of RNAs from this homology search to inhibit RNase L directly correlates to their E-value. The strong correlation between E-value and function is likely due to the sequence constraints of the 3C protease and the relatively small amount of phylogenetic variation in the sequences. Additionally, these functional ciRNAs all have at least 80% sequence identity to that of the PV1 ciRNA (Supplementary Fig. S4A&B). A phylogenetic tree made from VP1 protein sequence, a structural protein found upstream of the ciRNA’s genomic location, shows that the presence of a functional ciRNA tracks with the viral phylogeny (Fig. 3C).

**Figure 3.**
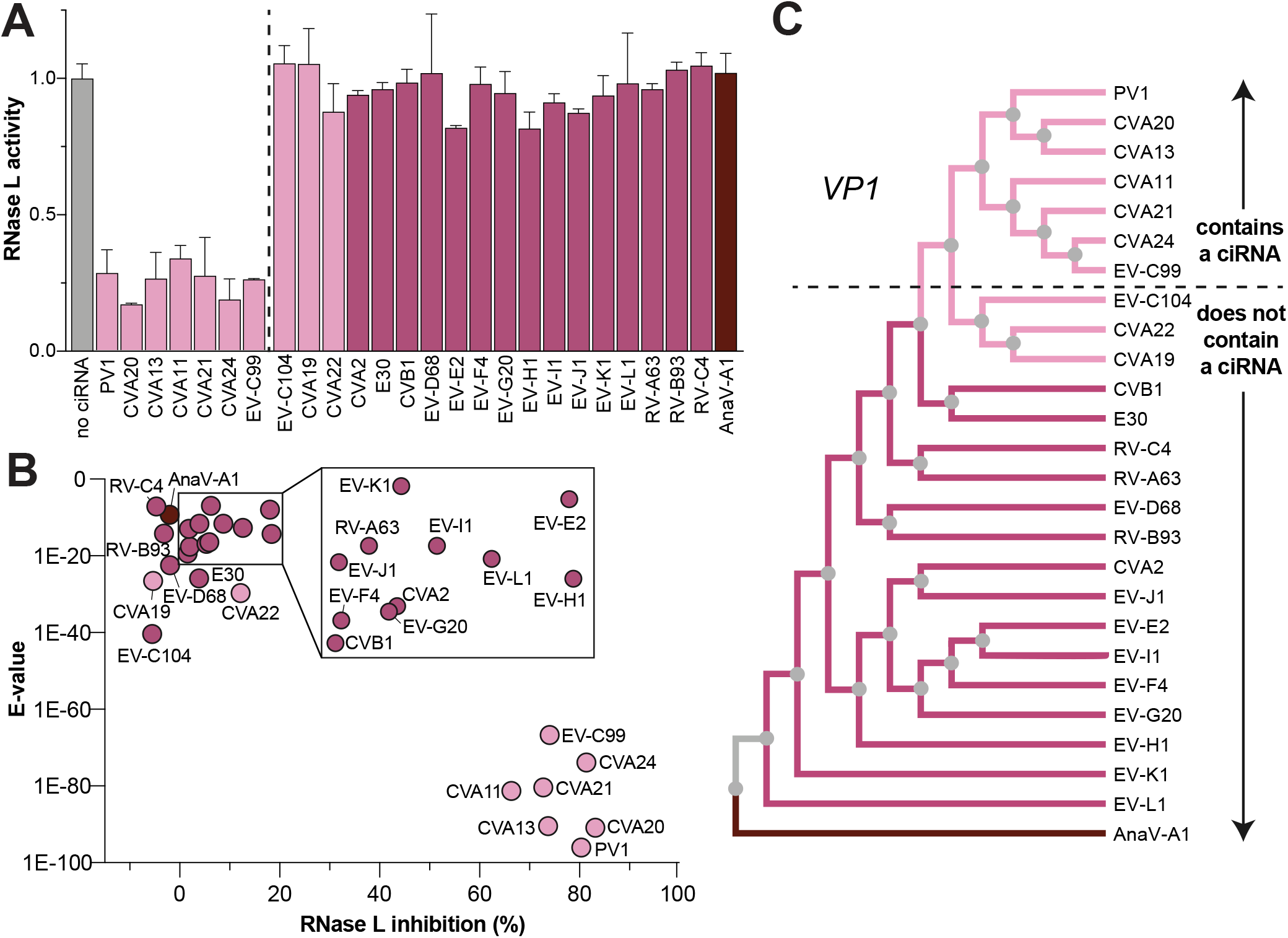
Functional ciRNAs are only found in Enterovirus coxsackiepol species. **(A)** Results of RNase L inhibition assay comparing RNAs from selected viral genomes (CV, coxsackievirus; EV, enterovirus; RV, rhinovirus; AnaV, anativirus). Reactions contained 100 nM ciRNA. Replicates (n=3) plotted at 40 min. **(B)** Percent inhibition of RNase L against their E-value. **(C)** Phylogenetic tree created using viral VP1 protein seqeunce. Dashed line separates viruses that contain a ciRNA from those that do not. **(A-C)** RNAs are colored according to their species: *Enterovirus coxsackiepol* (light pink), other *Enterovirus* species (magenta), and non-*Enterovirus* (dark red).

### Comparing functional and nonfunctional ciRNA secondary structures

Examination of the RNA sequences that emerged from the homology-based alignment but that do not inhibit RNase L revealed sequence differences in either the loop E motif, the L1-L4 interaction, or both. In the case of the sequence from CVA22 RNA, L4 is identical to that of the PV1 ciRNA but the CVA22 L1 sequence differs and, therefore, the kissing loop interaction cannot form (Fig. 4A). Additionally, this RNA has a loop E motif that differs from that found in PV1. This led us to speculate that the nonfunctional CVA22 RNA could be converted to a functional ciRNA just by changing these elements to match those found in PV1. To test this, we generated three sequences (Fig. 4A). The first, m11, replaces the CVA22 L1 sequence with one capable of base pairing with L4. The second, m12, replaces the degenerate loop E nucleotides with those from PV1, and the third combines these two mutations (m11+m12). At the same concentrations as the PV1 ciRNA (100 nM), neither the WT CVA22 RNA nor any of these mutants inhibit RNase L (Fig. 4B).

**Figure 4.**
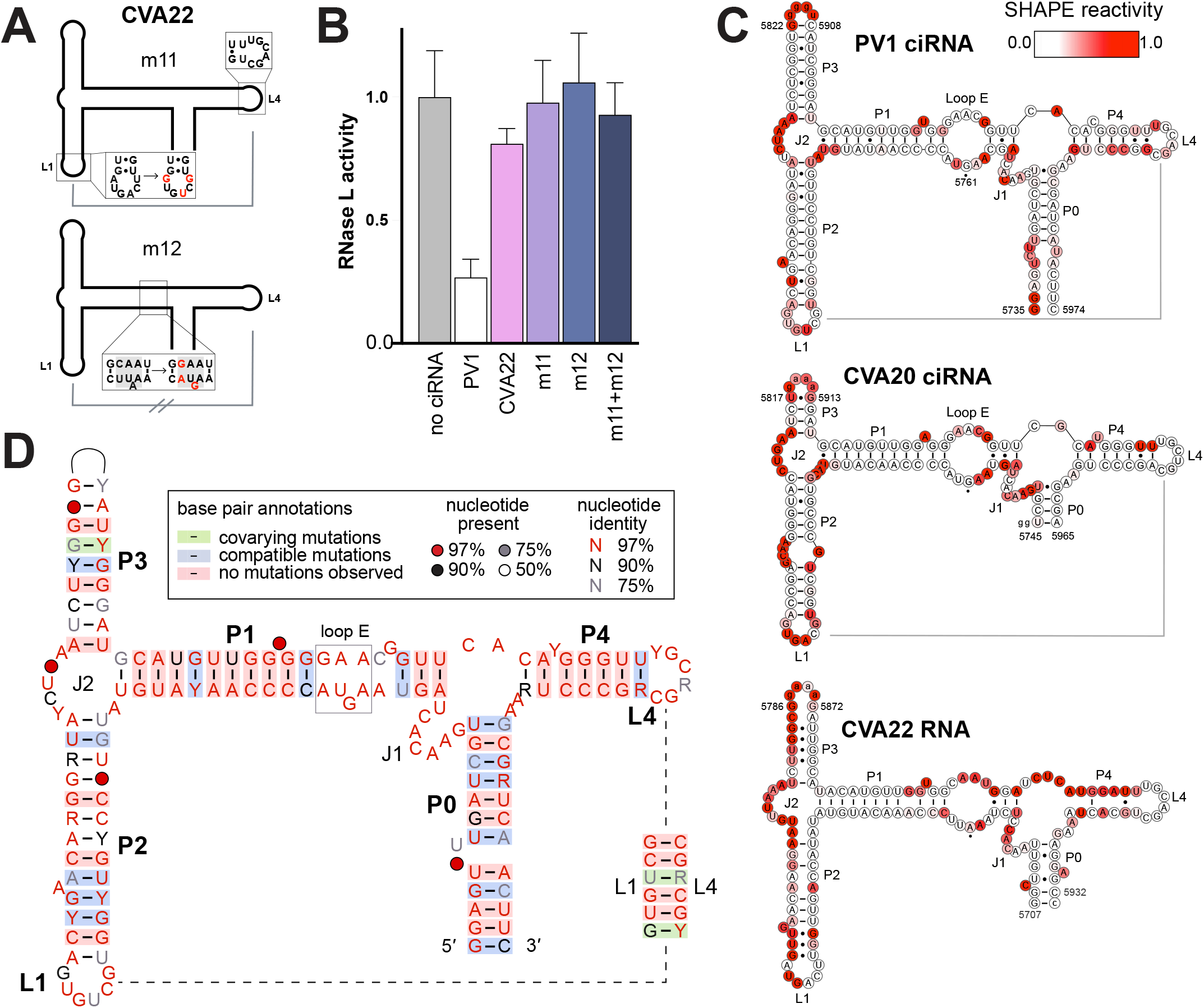
Secondary and tertiary structure requirements of a functional ciRNA vs a nonfunctional RNA. **(A)** Cartoon representation of mutations made to CVA22 RNA. Line between L1 and L4 represents the kissing loop. **(B)** RNase L inhibition assay of CVA22 RNA and its mutants at 100 nM RNA concentration. Replicates (n=3) plotted at 40 min. **(C)** Average SHAPE reactivities for PV1, CVA20, and CVA22 ciRNA (n=3 for each RNA). Nucleotide numbers correspond to their viral genome. Lower-case nucleotides correspond to non-endogenous sequences. **(D)** Covariation model of the ciRNA made in R2R. The interaction between L1 and L4 is represented with a 6-base pair stem.

The failure of these mutants to create a functional ciRNA from a nonfunctional version raises the question of why, even with the correct loop E and potential L1-L4 interactions in place, they do not inhibit RNase L. Does the CVA22 adopt a similar secondary structure to the PV1 ciRNA, or do differences in its fold limit function? To directly compare the secondary structures of functional PV1 ciRNAs with a nonfunctional but similar sequence, we performed selective 2′-hydroxyl acylation analyzed by primer extension and mutational profiling (SHAPE-MaP) on PV1 (functional), CVA20 (functional), and CVA22 (nonfunctional) RNAs.

The RNAs were subjected to chemical modification by 1M7, and reactivity values were plotted onto their predicted secondary structures (Fig. 4C). For the PV1 ciRNA, the data reveal low reactivity (<0.25) values for paired stems and high reactivity (>0.75) in loops, bulges, and junctions connecting these stems. Notably, the loops capping P3 has high reactivity, consistent with it not forming contacts with other regions of the ciRNA (Fig. 4C). Overall, the chemical probing pattern is consistent with the expected secondary structure. Likewise, the CVA20 RNA probing reactivity is consistent with a secondary structure homologous to the PV1 ciRNA, with minor differences in bulged nucleotides and internal loops (Fig. 4C). Interestingly, loops L1 and L4 for both CVA20 and PV1 show mild to moderate reactivity, suggesting that this tertiary interaction is somewhat transient, sampling a conformational ensemble, consistent with previous studies (Keel et al. 2012b). This experiment was performed in solution in the absence of RNase L; it possible that this kissing interaction is stabilized by RNase L binding.

In contrast to the functional CVA20 and PV1 ciRNAs, probing of the CVA22 RNA reveals a pattern that is inconsistent with a ciRNA-like secondary structure (Fig. 4C). Specifically, while the P0 and P1 stems of this RNA have low reactivity, indicating possible base pairing, the rest of the RNA is highly modified in stretches of nucleotides that would be protected if P2, P3, and P4 were stably forming. In fact, the secondary structure that is consistent with these data differs dramatically from that of the functional ciRNAs (Supplementary Fig. S5). These data explain why simply mutating the loop E motif and the kissing loop of CVA22 RNA was not sufficient to restore ciRNA function: the secondary structure does not provide the correct context for these necessary elements. These results indicate the high degree of structural specificity that underlies ciRNA’s ability to inhibit RNase L, requiring a combination of local and long-range structural elements within a precise structural architecture.

### Creating and using a consensus sequence and secondary structure model for functional ciRNA

Using the aligned sequences of all functional ciRNAs uncovered by our bioinformatic search, we created a consensus sequence and secondary structure model in R2R (Fig. 4D) (Weinberg and Breaker 2011). Most base pairs of the ciRNA do not covary, which is consistent with the very high degree of primary sequence conservation. However, many base pairs have compatible sequence variation, in which one side of the stem differs in sequence from PV1 but can form a viable base pair (e.g. A-U to G-U). Thus, many base pairs are completely conserved (Fig. 4D). It seems likely that many of these base pairs could be changed to other Watson-Crick pairs without a loss of function, but this has not naturally occurred due to the protein coding requirements of these RNAs. There are relatively few true compensatory changes, or regions where both bases mutate to maintain base pairing (e.g. C-G to U-A). Again, this is likely because the RNA sequence is limited by the constraints imposed by the 3C protease sequence.

Using the expanded alignment and information gleaned from our biochemical and structural characterization, we conducted additional homology-based searches using all fully sequenced, complete genomes in the RefSeq and GenBank viral genome databases (NCBI). Despite extensive searches, no new putative ciRNAs were found outside the *Enterovirus coxsackiepol* species, or that meet the E-value threshold that we established directly correlates with ciRNA function. To further expand our search, we exploited our discovery that the ciRNA could be circularly permuted. Specifically, we searched these databases using the circularly permuted sequence as an input to see if other viruses contained this same secondary structure in a different orientation. Again, no compelling examples were found. Thus, we conclude that functional PV-like ciRNAs only definitively exist in the *Enterovirus coxsackiepol* clade; if they exist in other RNA viruses, they must differ dramatically from the known architecture and sequence.

### Summary

Functional structured viral RNAs in UTRs are unconstrained by the need to encode a protein, which allows for the exploration of expansive sequence space, leading to extensive variation in sequence and structure. In contrast, ciRNAs must prioritize their role in encoding for the 3C protease protein, which constrains their evolution and likely limits diversity. It appears that ciRNAs have overcome this in part by forming the functional structure from noncontiguous sequence in the RNA genome, thus avoiding the most highly conserved sequences that encode the 3C protease active site. However, the examples of ciRNAs discovered in this work highlight how limited, unique, and highly specialized this type of ciRNA is across the viral world.

## Material and Methods

### RNA transcription and purification

Gene block fragments were ordered (Integrated DNA Technologies and TWIST Biosciences) with a 5′ T7 promoter sequence. Gene blocks were amplified by PCR and verified by 1% agarose gel electrophoresis. Transcription reactions were prepared with the following components: 30 mM Tris-Cl, pH 8, 7.5 mM each NTP, 30-60 mM MgCl2, 10 mM DTT, 0.1% spermidine, 0.1% Triton X-100, T7 polymerase and 25% (v/v) amplification product. Reactions were incubated overnight at 37°C and purified by denaturing 8% PAGE. RNA product was visualized by UV illumination and the RNA band corresponding to the correct RNA size was excised and then crushed using an RNase-free spatula. RNA was eluted from the crushed gel pieces in 20 mM sodium acetate, pH 5 overnight at 4°C. Eluted RNA was filtered (0.22 um, Millipore) and concentrated using 10-30 MWCO concentrators (depending on RNA size). RNA purity was assessed by denaturing 8% PAGE and RNA concentration was determined by Nanodrop.

### RNase L expression and purification

GST-RNase L fusion protein was expressed and purified as previously described (Dong et al. 1994). Expression vectors were a kind gift of the Silverman Lab at the Cleveland Clinic. Briefly, a pGEX expression vector containing wild type or mutant (H672N) RNase L (from homo sapiens, L10381.1/104-2329) was expressed in BL21(DE3) competent cells (New England Biolabs). RNase L protein expression was induced at OD_600_ = 0.8 using 0.3 mM IPTG at 18°C for 16-20 hr. Cells were harvested by centrifugation and pellets were resuspended in Buffer A (1X PBS, 10% glycerol, 1 mM EDTA, 5 mM MgCl2, 0.1 mM ATP, 14 mM BME) supplemented with 0.5 mg/mL lysozyme, 0.1 mg/mL DNase I, and 1 mM AEBSF. Cells were sonicated for 5 min at 50% amplitude and centrifuged for 30 min at 40k x g at 4°C. Supernatant was filtered using 0.22 um filter and incubated with 5 mL glutathione resin equilibrated in Buffer A for 2 hr at 4°C on a rocking platform. Resin was washed 3X with 10 CV Buffer A and washed 1X with 10 CV Buffer B (50 mM Tris pH 8.8, 100 mM KCl, 300 mM NaCl, 2 mM EDTA, 10 mM MgCl2, 0.2 mM ATP, 14 Mm BME). GST-RNase L fusion protein was eluted 3X in 1 CV of Buffer B supplemented with 20 mM reduced glutathione (pH 7.5). Elutions were pooled, concentrated and purified by size exclusion in 25 mM Tris, pH 7.5, 100 mM KCl, 10 mM MgCl2, 10% glycerol, 2 mM EDTA, 14 mM BME. Purity was assessed by size exclusion and by SDS-PAGE. Fractions were pooled, concentrated, flash-frozen in liquid nitrogen, and stored at −80°C.

### In vitro fluorescence-based RNase L activity assay

A 36-nt RNA probe (UA-rich sequence derived from RSV) containing a 5′ 6-FAM and 3′ BHQ1 (Sigma) was used as a substrate to measure RNase L activity (Thakur et al 2005). The 2′-5′-linked oligoadenylate (2-5A) was purchased as 5′-[ppp][rA2-5][rA2-5][rA]-3′ from ChemGenes. Each 50 uL experimental reaction contained 20 nM GST-RNase L, 20 nM 2-5A (for activated RNase L reactions), 25 mM Tris-Cl, pH 7.4, 100 mM KCl, 10 mM MgCl2, 50 µM ATP, 7 mM BME, and varying concentrations of ciRNA (0 nM – 1 µM). Fluorescence values (535 nm emission/485 nm excitation) were collected every minute for 1 hr at 25°C using a SpectraMax i3 Microplate Reader (Molecular Devices). Relative fluorescence units (RFUs) were averaged (n=3) and plotted against time. Standard deviations were calculated and plotted using GraphPad Prism.

### Alignment and homology search

The sequences for PV1, PV2, and PV3 ciRNA (NC_002058.3/5720-6023, M12197.1/5698-6173, and K01392.1/5720-6021, respectively) were aligned based on conserved secondary structure and used as a ‘seed’ to search for structure and sequence conservation in viral genomes (RefSeq and GenBank, Aug 2025) using INFERNAL (Eddy). This process was repeated using experimentally validated ciRNAs in the seed alignment.

### SHAPE-MaP chemical probing

SHAPE probing was performed as described in Smola et al. (Smola et al. 2015) RNA was denatured at 95°C for 3 min and cooled to room temperature and folded in 20 mM HEPES, pH 7.6, 100 mM NaCl, 10 mM MgCl_2_ at 37°C for 30 min. Five pmol of folded RNA (500 nM) was probed with 100 nmol 1M7 in DMSO (10 mM) or neat DMSO at 37°C for 75 s (5 half-lives). Reactions were quenched at 4°C for 5 min and purified using Monarch Kit (NEB), eluting in 14 µL RNase-free H_2_O. Modified and DMSO-treated RNA samples (3.5 pmol) were incubated with 0.8 µL 200 ng/µL random hexamers and 0.2 µL of 2 µM RNA-specific reverse primer at 65°C for 5 min and cooled on ice. Reactions containing 3.5 pmol RNA, 50 mM Tris, pH 8.0, 64 mM KCl, 12 mM MnCl2, 10 mM DTT, 0.5 mM each dNTP, 8 ng/µL random hexamers, 20 nM reverse primer, and 200 U SuperScript™ II reverse transcriptase (Invitrogen) were assembled at room temperature and incubated for 3 hr at 42°C. Reverse transcriptase was inactivated at 70°C for 15 min and reactions were purified with G-25 columns equilibrated in RNase-free H_2_O. Second strand synthesis was performed using the NEBNext® Ultra™ II Non-Directional RNA Second Strand Synthesis kit according to the manufacturer’s instructions apart from incubating the reaction for 2.5 hr at 16°C. Resulting dsDNA was purified using DNA Clean & Concentrator −5 kit (Zymo Research) and eluted in 10 µL RNase-free H_2_O. The concentration of each sample was determined using a Qubit 3 Fluorometer and Qubit dsDNA HS Assay Kit (Thermo). The sequencing library was prepared from 1 ng dsDNA using the Nextera XT DNA Library Preparation Kit (Illumina, FC-131-1024), following manufacturer’s protocol for DNA tagmentation and library amplification using Nextera XT Index Kit v2 Set A (Illumina) as index adapters. Library clean-up was performed using AMPure XP beads (Beckman Coulter) and concentration and quality of each sample was determined by Qubit and 4200 TapeStation System with a HS D5000 Screen Tape (Agilent). Samples were normalized, pooled, and were sequenced by a NovaSeq6000 (10 million paired reads per sample) at Novogene Corporation.

## Acknowledgements

Thank you to current and former Kieft Lab members for thoughtful discussion and technical assistance, including Katherine Segar for training in SHAPE-MaP probing and data analysis. This work was supported by NIH grants F31AI189061 (S.I.Z), R01AI133348 (J.S.K.). M.E.S was supported by a Jane Coffin Childs postdoctoral fellowship.

