## Supplemental Figures for "Enterovirus RNase L inhibiting RNAs are highly conserved with limited phylogenetic distribution"

**Supplemental information contents:**

**Supplemental Figure S1.** Graph of initial rates from in vitro inhibition assay

**Supplemental Figure S2.** Secondary structures of all mutants

**Supplemental Figure S3.** Alignment of putative ciRNA sequences

**Supplemental Figure S4.** List and graph of homology search E-value and percent identity to PV1

**Supplemental Figure S5.** Chemical probing of CVA22 putative ciRNA

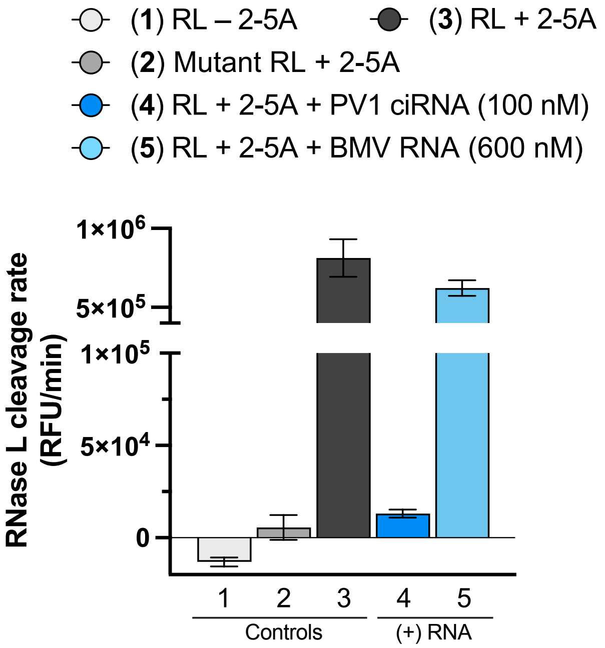


**Supplemental Figure S1. Graph of initial rates from in vitro inhibition assay.** An approximation of the initial rate of RNase L cleavage in the in vitro assay was obtained by measuring the slope of the fluorescence vs. time during the initial linear phase (RFU/min). Data are the same as those shown in Figure 1C&D.

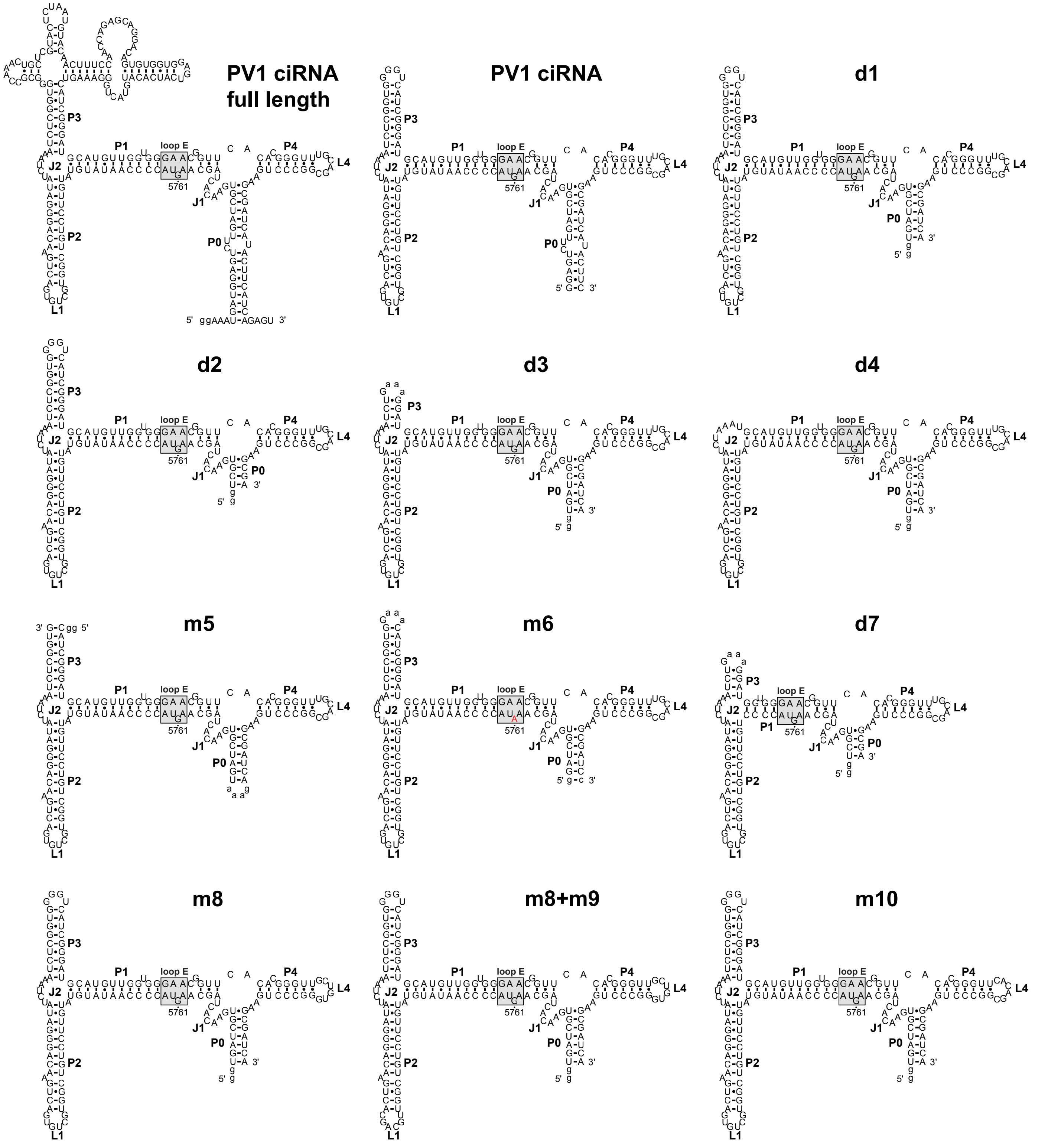


**Supplemental Figure S2. Secondary structures of all mutants.** The individual secondary structure of each mutant tested for RNase L inhibitory function. Results are shown in Figure 2B.

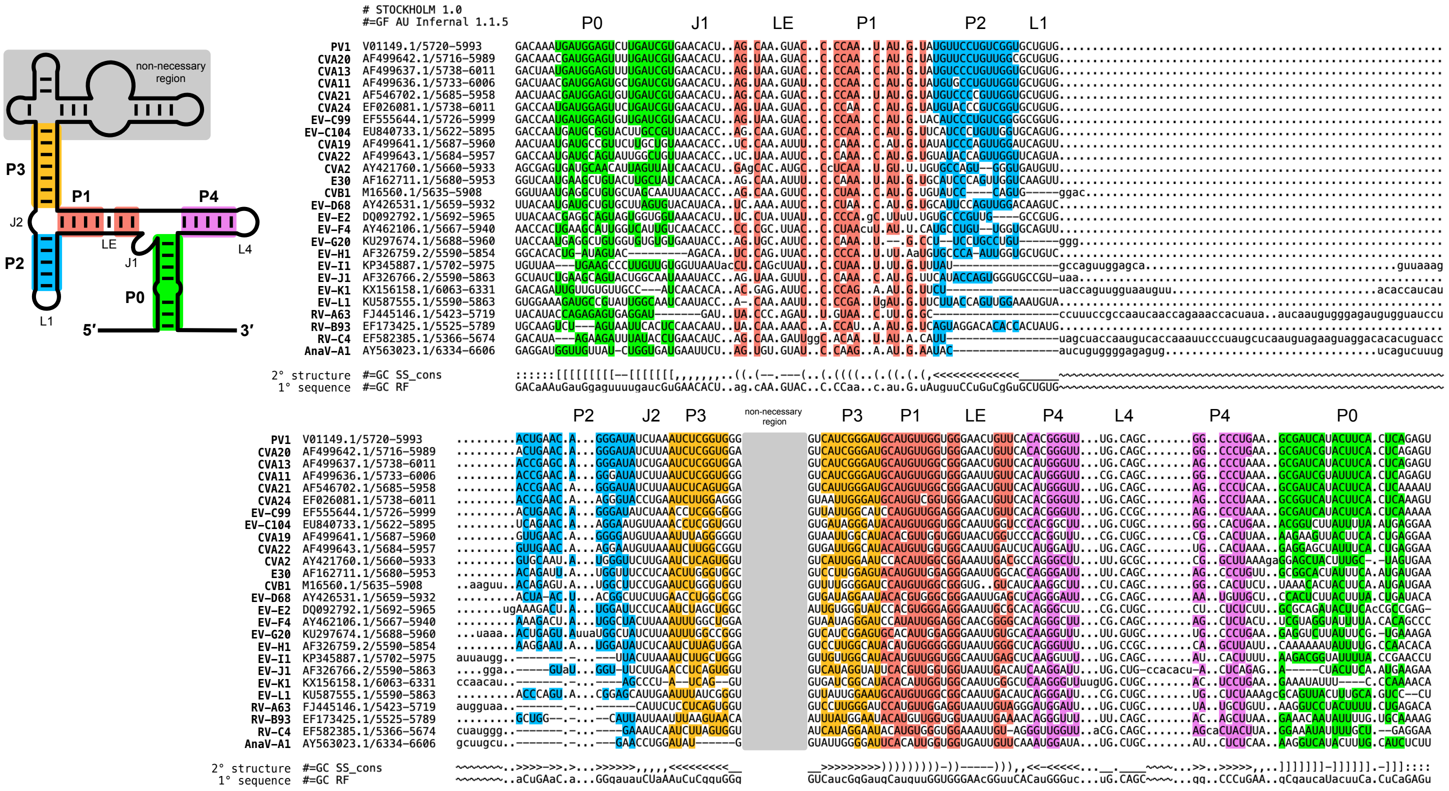


**Supplemental Figure S3. Alignment of putative ciRNA sequences.** Sequences of putative ciRNA sequences that came from the homology search are aligned and color coded by secondary structure element (left). Sequences are ranked by E-value.

**A**

| Virus | Genus | Species | Host | % identity (PV1) | E-value |
|---|---|---|---|---|---|
| PV1 | *Enterovirus* | *coxsackiepol* | human | 100 | 8.50E-97 |
| CVA20 | *Enterovirus* | *coxsackiepol* | human | 83.75 | 1.40E-91 |
| CVA13 | *Enterovirus* | *coxsackiepol* | human | 89.58 | 3.70E-91 |
| CVA11 | *Enterovirus* | *coxsackiepol* | human | 90.42 | 4.80E-82 |
| CVA21 | *Enterovirus* | *coxsackiepol* | human | 90 | 4.60E-81 |
| CVA24 | *Enterovirus* | *coxsackiepol* | human | 88.33 | 1.40E-74 |
| EV-C99 | *Enterovirus* | *coxsackiepol* | human | 90.83 | 1.80E-67 |
| EV-C104 | *Enterovirus* | *coxsackiepol* | human | 72.5 | 4.00E-41 |
| CVA19 | *Enterovirus* | *coxsackiepol* | human | 67.5 | 3.10E-27 |
| CVA22 | *Enterovirus* | *coxsackiepol* | human | 68.33 | 2.10E-30 |
| CVA2 | *Enterovirus* | *alphacoxsackie* | human | 60.42 | 3.60E-17 |
| E30 | *Enterovirus* | *betacoxsackie* | human | 47.5 | 1.40E-26 |
| CVB1 | *Enterovirus* | *betacoxsackie* | human | 65.42 | 4.20E-20 |
| EV-D68 | *Enterovirus* | *deconjunct* | human | 46.67 | 3.30E-23 |
| EV-E2 | *Enterovirus* | *eibovi* | bovine | 46.25 | 1.10E-08 |
| EV-F4 | *Enterovirus* | *fituari* | possum | 64.58 | 2.60E-18 |
| EV-G20 | *Enterovirus* | *geswini* | goat | 48.52 | 1.30E-17 |
| EV-H1 | *Enterovirus* | *hesimi* | ape | 43.33 | 5.40E-15 |
| EV-I1 | *Enterovirus* | *idromi* | camel | 44.58 | 2.20E-12 |
| EV-J1 | *Enterovirus* | *jesimi* | ape | 46.25 | 1.10E-13 |
| EV-K1 | *Enterovirus* | *krodeni* | rodent | 45.69 | 1.20E-07 |
| EV-L1 | *Enterovirus* | *lesimi* | ape | 44.58 | 2.00E-13 |
| RV-A63 | *Enterovirus* | *alpharhino* | human | 42.55 | 2.20E-12 |
| RV-B93 | *Enterovirus* | *betarhino* | human | 43.1 | 6.30E-15 |
| RV-C4 | *Enterovirus* | *cerhino* | human | 44.87 | 7.10E-08 |
| AnaV-A1 | *Anativirus* | *aductai* | duck | 57.92 | 5.40E-10 |

**B**

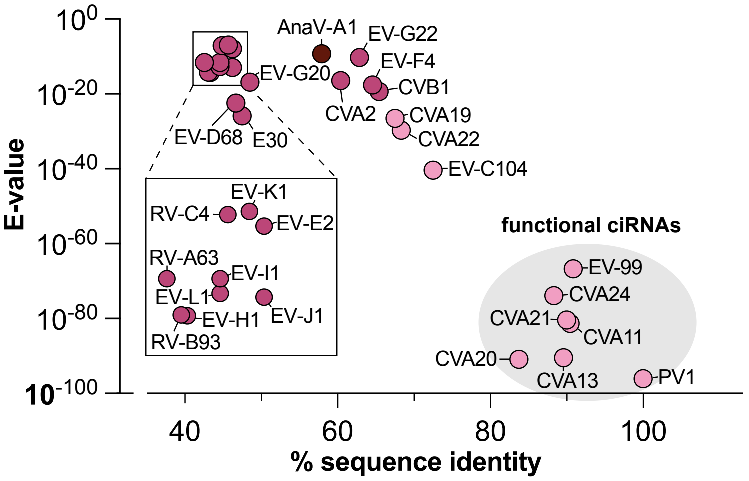


**Supplemental Figure S4. List and graph of homology search E-value and percent identity to PV1. A.** List of the putative ciRNA sequences that were identified in our bioinformatic search, with virus in which they are found, and the host species. Percent identity and E-value are shown. **B.** Graph of E-value v. percent sequence identity to PV1. RNAs that we identified as functional ciRNAs are clustered and indicated with a shaded oval.

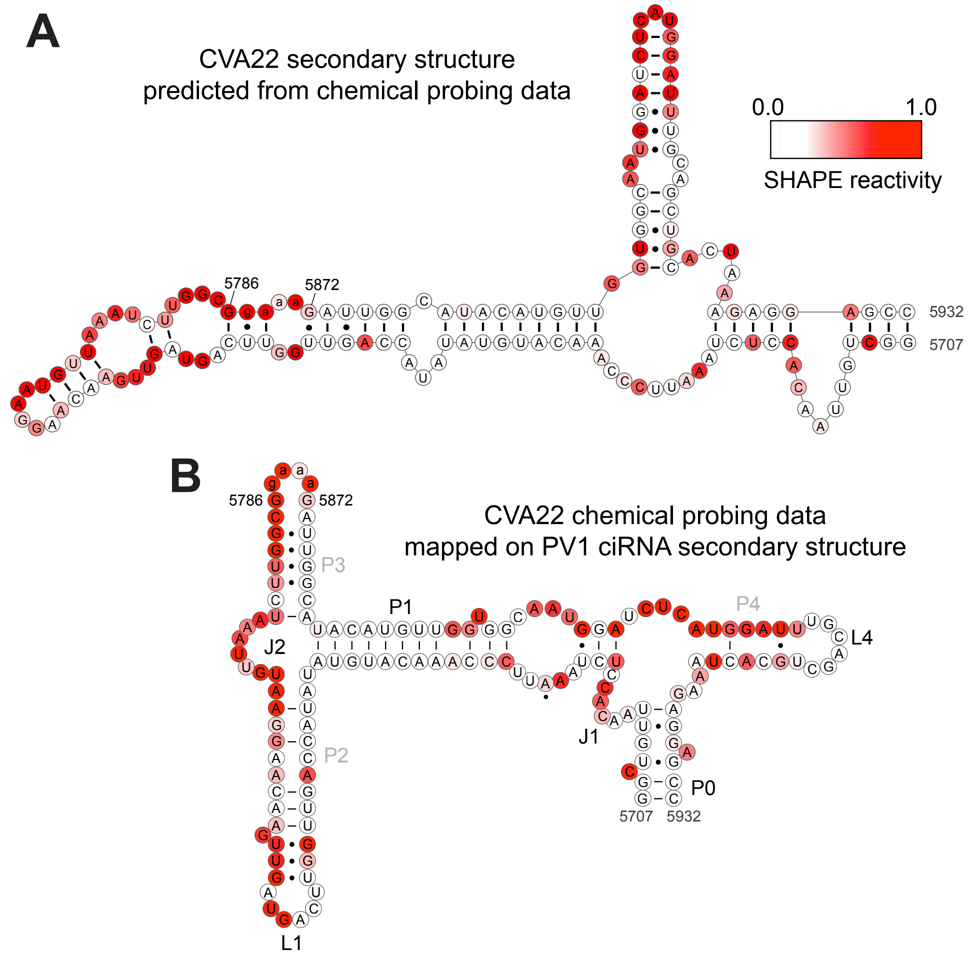


**Supplemental Figure S5. Chemical probing of CVA22 putative ciRNA. A.** Chemical reactivity from SHAPE probing mapped onto the predicted secondary structure of the non-inhibitory CVA22 sequence identified in our search. The predicted secondary structure is a result of analysis of the probing and does not match the known secondary structure of the inhibitory ciRNA from PV1 (and others). **B.** CVA22 depicted in the secondary structure from the PV1 ciRNA, with the CVA22 chemical probing data mapped onto the structure. The chemical probing pattern does not match the structure. The conclusion from these data is that the CVA22 RNA does not adopt a PV1 ciRNA-like secondary structure.
